# The complement regulator CD55 modulates TLR9 signaling and supports survival in marginal zone B cells

**DOI:** 10.1101/2024.03.01.582833

**Authors:** Iris Lee, Ivana Ling, Sonam Verma, Wumei Blanche, Christine T.N. Pham, Peggy L. Kendall

## Abstract

Marginal zone (MZ) B cells bridge innate and adaptive immunity by sensing bloodborne antigens and producing rapid antibody and cytokine responses. CD55 is a membrane-bound complement regulator that interferes with complement activation, an important component of innate immunity. CD55 also regulates adaptive immunity—CD55 downregulation is critical for germinal center reactions. MZ B cells also express low CD55, but its role in MZ B cell function is unknown. Using germline knockout mice, we found that similar numbers of MZ B cells are initially established in 3-week-old CD55-deficient mice compared to wild-type (WT) mice. However, MZ B cells fail to accumulate as mice age and undergo increased apoptosis. Following *ex vivo* stimulation of MZ B cells through Toll-like receptor 9, we observed a proinflammatory phenotype with increased IL-6 expression. These findings demonstrate a critical role for CD55 in supporting MZ B cell survival while also regulating cellular function.

## Introduction

Marginal zone (MZ) B cells are an innate-like lymphocyte population that produce rapid responses against antigens in the blood. Upon encountering antigens, MZ B cells rapidly differentiate into plasmablasts and produce T cell-independent, low affinity, polyreactive immunoglobulin M (IgM) responses and cytokines.^1, 2^ In addition to foreign antigens, a high proportion of their B cell receptors also recognize self-antigen,^3, 4^ which serve housekeeping purposes for clearing self-antigen but may also contribute to loss of tolerance when dysregulated.^4, 5^

The complement system, known for its canonical roles in innate defense against pathogens, provides complex regulation of B cells in the adaptive immune response. MZ B cells use complement receptors 1 and 2 (CD35 and CD21, respectively) to bind and transport complement-opsonized antigens to follicular dendritic cells and follicular B cells.^6, 7^ Complement-opsonized antigens bound to CD21 and CD35 in turn mediate B cell activation and proliferation.^8, 9^ More recent studies found that complement fragments signaling through C3aR1 and C5aR1 also directly impact B cell function. For example, in response to T cell-dependent antigens, C3aR1 and C5aR1 signaling mediate B cell activation, proliferation, antibody production, and class switch recombination.^10, 11^

Complement regulators are crucial for fine-tuning the activation of complement and have newly defined roles in humoral immunity. The membrane-anchored complement regulator CD55 has both complement-independent and complement-dependent modes of action. CD55 binds to CD97, an adhesion G-protein-coupled receptor; this interaction aids in cellular homing.^12, 13^ It also interferes with the assembly of C3 convertase, limiting complement activation. After stimulation, follicular B cells entering the germinal center downregulate CD55 to allow C3 cleavage, production of complement fragments, and activation of C3aR1 and C5aR1 to facilitate positive selection within the germinal center.^14, 15^ CD55 is then upregulated as germinal center B cells further differentiate into memory B cells or plasma cells.^14, 15^

Immature B cells also modulate CD55 expression. In human bone marrow, immature B cells express low amounts of CD55, which is upregulated as the cells leave the bone marrow and become transitional B cells.^14^ While MZ B cells and follicular B cells both develop from transitional B cells,^16, 17^ MZ B cells express less CD55 than follicular B cells.^18^ Recently, the interaction between CD97 on MZ B cells and CD55 on erythrocytes was found to be important for homing of MZ B cells to their splenic niche.^18^ However, little is known about the role of CD55 in the development or biology of MZ B cells.

MZ B cells are expanded in multiple mouse models of autoimmunity as well as Sjogren’s syndrome and Graves’ disease in humans.^19, 20^ In particular, MZ B cells play a pivotal role in the initiation of disease in the collagen-induced arthritis (CIA) mouse model of rheumatoid arthritis. Following subdermal injection of type II collagen, MZ B cells expand rapidly, produce anti-collagen antibodies and cytokines, and have significant antigen-presenting capability to cognate T cells.^4^ Loss of CD55 in this model delayed onset of arthritis and attenuated disease, but MZ B cells were not examined. Interestingly, loss of CD97 also delayed onset of arthritis but did not attenuate disease severity suggesting that CD55 has homing-independent functions in CIA. Thus, we hypothesized that loss of CD55 would alter MZ B cell function. We found that loss of CD55 increased apoptosis in MZ B cells at homeostasis but increased IL-6 expression after stimulation with a TLR9 agonist. These findings provide new insights regarding the influence of complement on MZ B cells.

## Materials and Methods

### Mice

C57BL/6 CD55 KO mice were a kind gift from Wen-Chao Song (University of Pennsylvania, Philadelphia, PA). C57BL/6 mice were originally obtained from The Jackson Laboratory (Bar Harbor, ME) and subsequently bred and maintained in our mouse facility. 9-12-week-old age- and sex-matched male and female mice were used for all experiments except where specified. Mice were maintained in the same facility and were littermates where described. All studies are approved by the Washington University Institutional Animal Care and Use Committee (protocol 22-0341).

### Flow cytometry and antibodies

Single-cell suspensions were obtained and stained using fluorochromes against B220 (RA3-6B2), IgMb (AF6-78 or eB121-15F9), IgD (11-26c.2a), CD21 (7G6), CD23 (B3B4), and CD55 (R&D Systems 583905). LIVE/DEAD Fixable Blue or Violet Stain (ThermoFisher) excluded dead cells. Intracellular staining was conducted using the FoxP3/Transcription Factor Staining kit (eBioscience) per manufacturer’s instructions. Cell survival stains were conducted with Bcl-xL (54H6; Cell Signaling) and cytokine stains with IL-6 (MP5-20F3; Biolegend), IL-10 (JES5-16E3; Biolegend), and TNFα (MP6-XT22; Biolegend). Bromodeoxuridine (BrdU) incorporation was performed per the manufacturer’s instructions (BD Pharmingen). Activated caspase-3/7 staining was performed per manufacturer’s instructions (Immunochemistory Technologies). Samples were analyzed using a Cytek Aurora spectral flow cytometer (Cytek Biosciences), and data was analyzed using FlowJo software (version 10.9.0; Tree Star).

### Histology

Frozen spleens were fixed using a 1:1 ratio of chilled methanol for 10 minutes at room temperature followed by two washes with phosphate buffered saline (PBS). After blocking with 2% bovine serum albumin (BSA) in PBS, tissues were incubated with unconjugated anti-rabbit CD19 (Abcam # ab245235) and anti-rat MOMA1 (Abcam #ab53443) primary antibodies overnight at 4°C followed by incubation with anti-rabbit Alexa 488 and anti-rat Alexa 555 secondary antibodies in 2% BSA. Counterstaining was performed using Hoechst dye (ThermoFisher) for 10 minutes at room temperature and fixation in ProLong Diamond Antifade Mountant (ThermoFisher). Images were captured from non-overlapping areas using a Zeiss LSM 880 with Airyscan Confocal Microscope (Carl Zeiss, Germany).

### Transcriptome analysis

B cells from 14-week-old mice were negatively selected after incubation of splenocytes with anti-CD11b-biotin, anti-CD11c-biotin, and anti-CD43-biotin (BD Biosciences) followed by streptavidin beads (Miltenyi) as described previously.^21^ Lymphocytes were then purified using MACS LS Columns per the manufacturer’s instructions and sorted on the iCyt Synergy (Sony) into Buffer RLT (Qiagen). RNA was next extracted using the RNeasy Micro Kit (Qiagen) and submitted for library preparation and sequencing to the Genome Technology Access Center at the McDonnell Genome Institute at Washington University in St. Louis. Briefly, samples were prepared according to the SMARTer Ultra Low RNA kit for Illumina Sequencing (Takara-Clontech) per manufacturer’s protocol, indexed, pooled, and sequenced on an Illumina NovaSeq 6000. Base calls and demultiplexing were performed with Illumina’s bcl2fastq2 software. RNA sequencing (RNA-Seq) reads were then aligned and quantitated to the Ensembl release 101 primary assembly with an Illumina DRAGEN Bio-IT on-premise server running version 3.9.3-8 software.

All gene counts were imported into the R/Bioconductor package EdgeR^22^ and TMM normalization size factors were calculated to adjust samples for differences in library size. Ribosomal genes and genes not expressed in the smallest group size minus one samples greater than one count-per-million were excluded from further analysis. The TMM size factors and the matrix of counts were then imported into the R/Bioconductor package Limma.^23^ Weighted likelihoods based on the observed mean-variance relationship of every gene and sample were calculated for all samples and the count matrix was transformed to moderated log 2 counts-per-million with Limma’s voomWithQualityWeights.^24^ The performance of all genes was assessed with plots of the residual standard deviation of every gene to their average log-count with a robustly fitted trend line of the residuals. Differential expression analysis was then performed to analyze for differences between conditions and the results were filtered for only those genes with Benjamini-Hochberg false-discovery rate adjusted p-values less than or equal to 0.05. Heatmaps were generated using the pheatmap package in RStudio (v2023.12.0+369, Posit). Clustering was performed using Euclidian.

We imputed normalized gene expression data from our RNA-Seq experiment into gene set enrichment analysis (GSEA) software (4.1.0, Broad Institute). GSEA then analyzed our dataset for enriched genetic signatures curated in the hallmark gene sets by the Molecular Signatures Database.^25, 26^ Genes were ranked based on their expression and compared against the hallmark gene sets to generate an enrichment score. A nominal p-value was then generated followed by normalization for the size of the gene set and adjustment for multiple hypothesis testing to yield a false discovery rate (FDR).^26^ Gene sets were considered significant with a nominal p-value < 0.01 and FDR q-val < 0.1. Differentially expressed genes with Benjamini-Hochberg FDR adjusted p-values <= 0.05 were also examined using a knowledge engine called COMPBIO.^27^

### B cell activation

Splenocytes were cultured in complete RPMI with 2% heat inactivated fetal bovine serum (Gibco) for activation assays at 4 ×10^6^ cells/mL for cell survival or 10×10^6^ cells/mL for cytokine production. Cells were unstimulated or stimulated by incubating with 10 μg/mL goat anti-mouse IgM (Jackson ImmunoResearch) or 1 μM CpG (ODN 1826 (TLRGRADE®); Enzo).

### Statistical analysis

Statistical significance was determined using Student’s t-test, with Welch’s correction used when variances were unequal, or two-way ANOVA followed by Šídák’s multiple comparisons test. P-values were corrected for multiple comparisons using Holm-Šídák’s correction. Statistical calculations were performed using GraphPad Prism 10.0.3 for macOS (GraphPad Software, La Jolla, CA).

## Results and Discussion

Given that MZ B cells are involved in the early pathogenesis of CIA and are expanded in multiple models of autoimmunity, we sought to understand the importance of CD55 to MZ B cell biology. We first used flow cytometry to evaluate CD55 surface expression levels across splenic B cell populations in WT mice (**Fig. 1A**). We found relatively low CD55 expression in early transitional (T1) and MZ populations and high CD55 expression in late transitional (T2) and follicular B cell populations.

**Figure 1.**
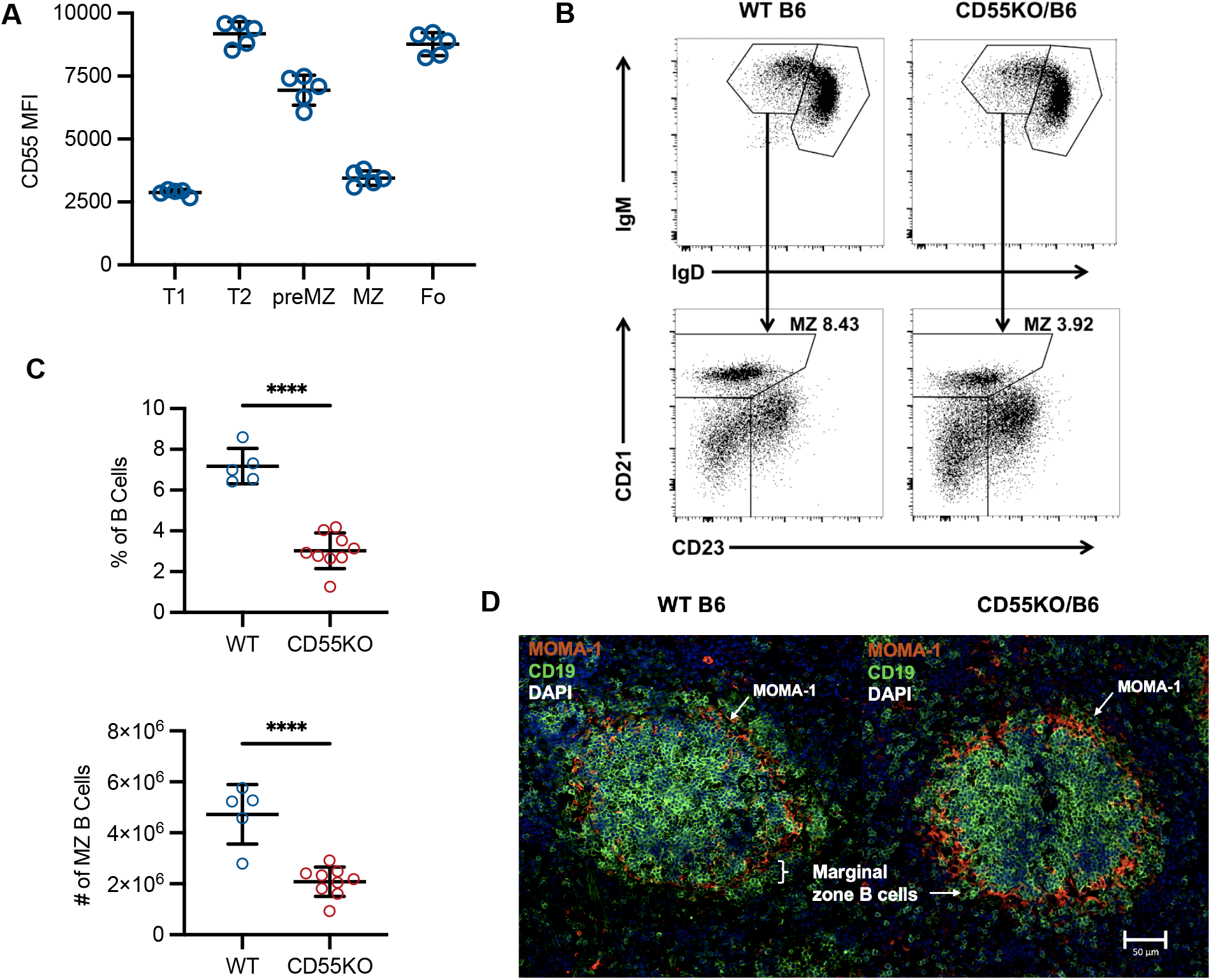
Loss of CD55 decreases MZ B cell subsets within the spleen. **A** CD55 expression is lower on MZ B cells compared to transitional 2 (T2), preMZ, and follicular (Fo) B cells. **B** Flow cytometry gating of MZ B cells in WT (left) and CD55 KO (right) mice. **C** Percentage and number of MZ B cells in 10- to 12-week-old WT and CD55 KO mice. Statistical significance determined using Student’s t-test. Error bars represent mean +/- standard deviation. ****p≤0.0001 **D** Immunofluorescence staining of frozen spleen sections showing a normal and attenuated MZ B cell zone in 12-week-old male WT (left) and CD55 KO (right) mice, respectively. Sections were stained for B cells (CD19) and metallophilic macrophages (MOMA-1). White bracket and arrows indicate MZ B cells located outside of the metallophilic macrophage ring.

We next analyzed splenocytes isolated from WT and CD55 KO mice using flow cytometry and observed that CD55 KO mice had lower frequencies and total numbers of MZ B cells compared to WT at 10-12 weeks of age (**Fig. 1B-C**). Similarly, immunofluorescence microscopy of splenic sections showed decreased MZ B cells in CD55 KO mice relative to WT (**Fig. 1D**) similar to the results of Liu et al.^18^ In contrast, the percentage and number of follicular B cells was unchanged between WT and CD55 KO mice (**Fig. 2B-C**).

**Figure 2.**
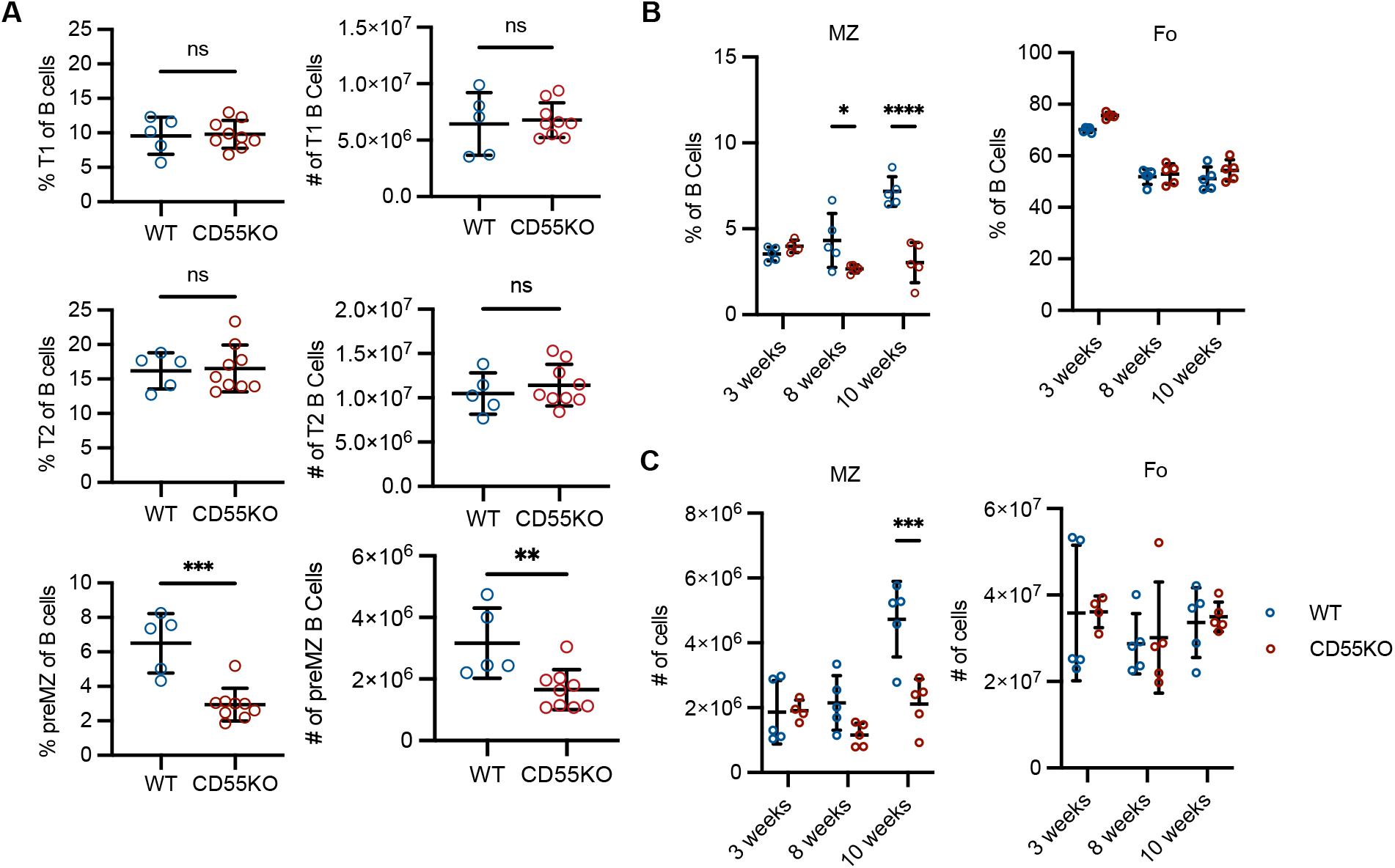
CD55 is dispensible for Follicular and MZ B cell development but is necessary for MZ B cell expansion over time. **A** Percentage (left) and absolute number (right) of transitional 1 (T1), transitional 2 (T2), and preMZ B cells in 10- to 12-week-old WT and CD55 KO mice. **B** Percentage and **C** number of MZ and follicular (Fo) B cells in 3-, 8-, and 10-week-old WT and CD55 KO mice. **A-C** Statistical significance determined using Student’s t-test (**A**) or two-way ANOVA (**B-C**) adjusted for multiplicity using Holm-Šídák’s correction (**B-C**). Error bars represent mean +/- standard deviation. ns not significant; *p≤0.05; **p≤0.01; ***p≤0.001; ****p≤0.0001.

Since MZ B cells were decreased in spleens of CD55 KO mice, we then evaluated the developmental precursors of MZ B cells. Using flow cytometry, we found no differences between early transitional (T1: IgM^high^, IgD^low^, CD23^low^, CD21/35^low^) or late transitional (T2: IgM^high^, IgD^high^, CD23^high^, CD21/35^mid^) B cells. There was no accumulation of transitional cells that would indicate failure of B cells at these early stages to mature further when CD55 is absent. Premarginal zone B cells (IgM^high^, IgD^high^, CD21^high^), the direct precursors of MZ B cells, were reduced in CD55 KO mice, indicating that CD55 contributions to MZ B cell development begins at this stage (**Fig. 2A**).

The MZ B cell population is initially established at three weeks of age in WT mice. These long-lived B cells then accumulate over time. To examine potential CD55 contributions to development, we next compared three-week-old, eight-week-old, and ten-week-old WT and CD55 KO mice. At three weeks of age, there were no differences in the numbers of premarginal zone or MZ B cells. Both WT and CD55 KO mice had approximately 2×10^6^ MZ B cells indicating that CD55 is not required for the MZ population to be established in the spleen. At eight weeks of age, there was a significant difference in the percentage of MZ B cells without a significant difference in the number of MZ B cells **(Fig. 2B-C**). Between eight and ten weeks of age, we observed that the percentage and number of MZ B cells in WT, but not CD55 KO mice, increased significantly (4.7 × 10^6^ vs 2.1 × 10^6^; p<0.0001) (**Fig. 2B-C**). The differences noted in ten week old mice were preserved at 17 weeks of age (**Supp Fig. 1**). In contrast, the percentage and number of follicular B cells was unchanged between WT and CD55 KO mice at any age (**Fig. 2B-C)**.

To understand the basis for these findings, we performed bulk RNA-Seq on flow-sorted MZ B cells from 14-week-old CD55 KO and WT mice. RNA-Seq analysis showed decreased transcripts related to chaperones of heat shock proteins (*STIP1, AHSA1, FKBP4, HSP90b1*, and *DNAjb11*) (**Fig. 3A)**. Heat shock proteins are upregulated during environmental stresses and inhibit cell death signaling. Pathway analysis using the hallmark library of gene signatures showed that CD55 regulates pathways related to cell cycle, survival, and death. Genes in the G2M (p<0.001; FDR q-val = 0.004), E2F (p<0.001; FDR q-val = 0.006), and MTORC1 pathways (p<0.001; FDR q-val = 0.059) were depleted in CD55 KO MZ B cells (**Fig. 3C)**. Further analysis of cell pathways with CompBio (Comprehensive Multi-omics Platform for Biological InterpretatiOn), an ontology-free approach, similarly showed HSP90 chaperone cycle and negative regulation of apoptosis as major themes that are downregulated in CD55 KO MZ B cells **(Fig. 3B)**.

**Figure 3.**
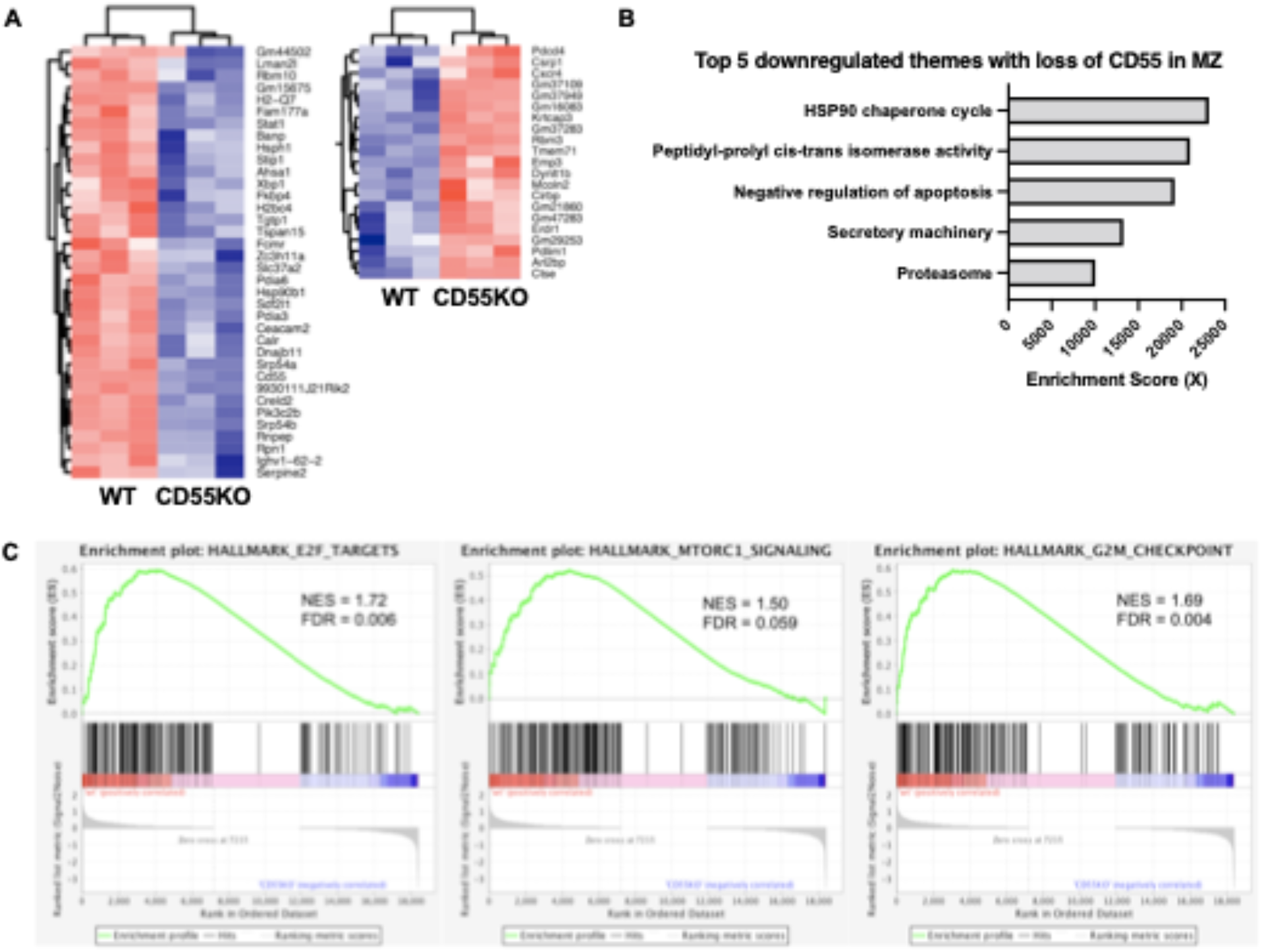
CD55 regulates cell cycle and apoptotic pathways. **A** Bulk RNA-Seq of MZ B cells sorted as B220^high^,CD23^low^,CD21/35^high^ from 14-week-old male WT (n=3) and CD55 KO (n=3) mice. Top genes from differential expression analysis were filtered for Benjamini-Hochberg false-discovery rate adjusted p≤0.05. Mice clustered together by genotype. **B** Top five significant themes from COMPBIO generated from RNA-Seq data plotted by their enrichment scores. **C** Significantly enriched (nominal p<0.01 and FDR q< 0.1) hallmark gene pathways using GSEA of RNA-Seq data showing downregulation of cell cycle (E2F Targets, G2M Checkpoint) MTORC1 Signaling pathways.

Given that RNA-Seq identified cell cycle control and apoptosis as key differences between CD55 KO and WT MZ B cells, we probed the ability of MZ B cells to proliferate or undergo apoptosis at homeostasis. We injected bromodeoxyuridine (BrdU) into WT and CD55 KO mice and harvested splenocytes 24 hours later. MZ B cells in CD55 KO mice showed identical BrdU incorporation to WT suggesting that CD55 is dispensable for MZ proliferation at homeostasis (**Fig. 4**). In contrast, caspase staining showed an increase in the percentage of MZ B cells with low levels of activated caspase-3 and caspase-7 in CD55 KO MZ B cells, indicative of increased apoptosis in the absence of CD55 (**Fig. 4**). Since MZ B cells are exposed to blood-borne antigens in the marginal zone of the spleen, we next probed the ability of MZ B cells to regulate prosurvival signals in response to stimulation. Loss of CD55 resulted in slower upregulation of the prosurvival factor Bcl-xL in response to CpG stimulation with significantly lower Bcl-xL expression at 4 hours, but equal Bcl-xL expression at 7 hours *ex vivo* (**Fig. 4**). Follicular B cells did not have a difference in the percentage of cells expressing activated caspase-3 and caspase-7 with loss of CD55, but also delayed upregulation of Bcl-xL in response to CpG stimulation. These findings suggest that loss of CD55 causes an increase in MZ cell death through inadequate or delayed expression of prosurvival factors potentially through loss of the CD97-CD55 interaction^18^ or stimulation by PAMPs. While follicular B cells also delayed upregulation of Bcl-xL in response to stimulation through TLR9, they are not exposed directly to antigens in the blood, unlike MZ B cells.^6^

**Figure 4.**
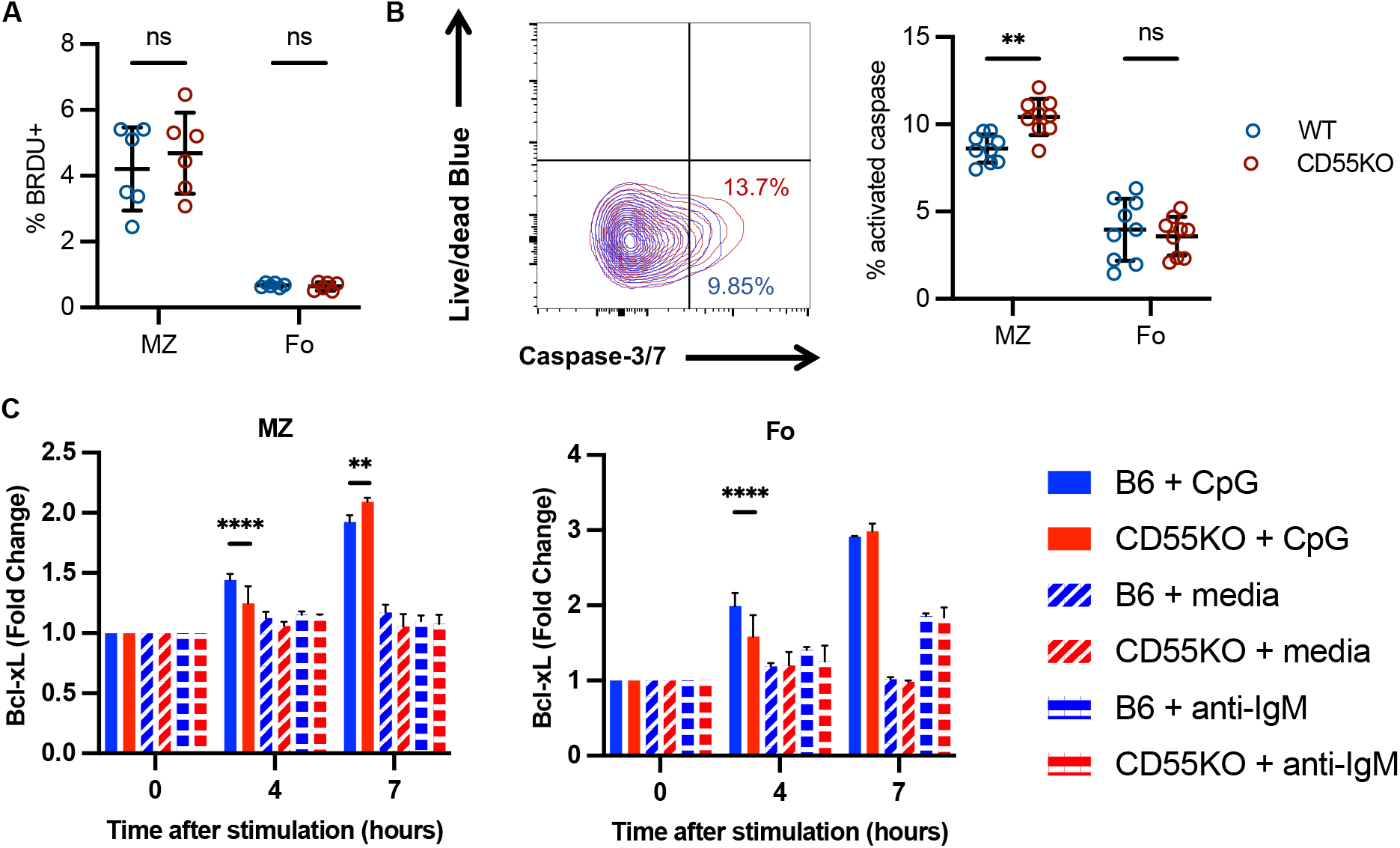
Loss of CD55 causes increased cell death in MZ B cells without loss of proliferative capacity *in vivo*. **A** Percentage of MZ and follicular (Fo) B cells following 24 hours of *in vivo* BrdU incorporation (via intraperitoneal injection). **B** Percentage of MZ and follicular B cells with activated caspase-3 and -7 expression including littermates. **C** Fold-change of Bcl-xL in WT (blue) and CD55 KO (red) MZ and follicular B cells cultured with CpG (solid lines), media alone (dashed lines), or anti-IgM (dotted lines) including littermates. **A-C** Statistical significance determined using t-test (**A-B**) or or two-way ANOVA (**C**) adjusted for multiplicity using Holm-Šídák’s correction. Error bars represent mean +/- standard deviation. ns not significant; **p≤0.01; ****p≤0.0001.

MZ B cells respond to bloodborne antigens by rapidly producing antibodies and cytokines. So next we examined serum antibody levels and cytokine production after stimulation. Baseline serum IgM expression was higher in CD55 KO compared to WT mice (**Fig. 5A**). In addition, a higher percentage of CD55 KO MZ B cells expressed IL-6 after CpG stimulation (**Fig. 5B)**. There were no differences between CD55 KO and WT MZ B cells for IL-10 or TNFα expression (**Fig. 5C-D**). There were also no differences between CD55 KO and WT follicular B cells for IL-6, IL-10, or TNFα expression. Since loss of the complement receptor CD21 increased MZ IL-6 production in a mouse model of arthritis,^4^ we next examined the expression of complement receptors on B cells. CD21/35 was decreased in all CD55 KO B cells with the greatest magnitude difference in CD55 KO MZ B cells compared to WT (**Supp Fig. 2**).

**Figure 5.**
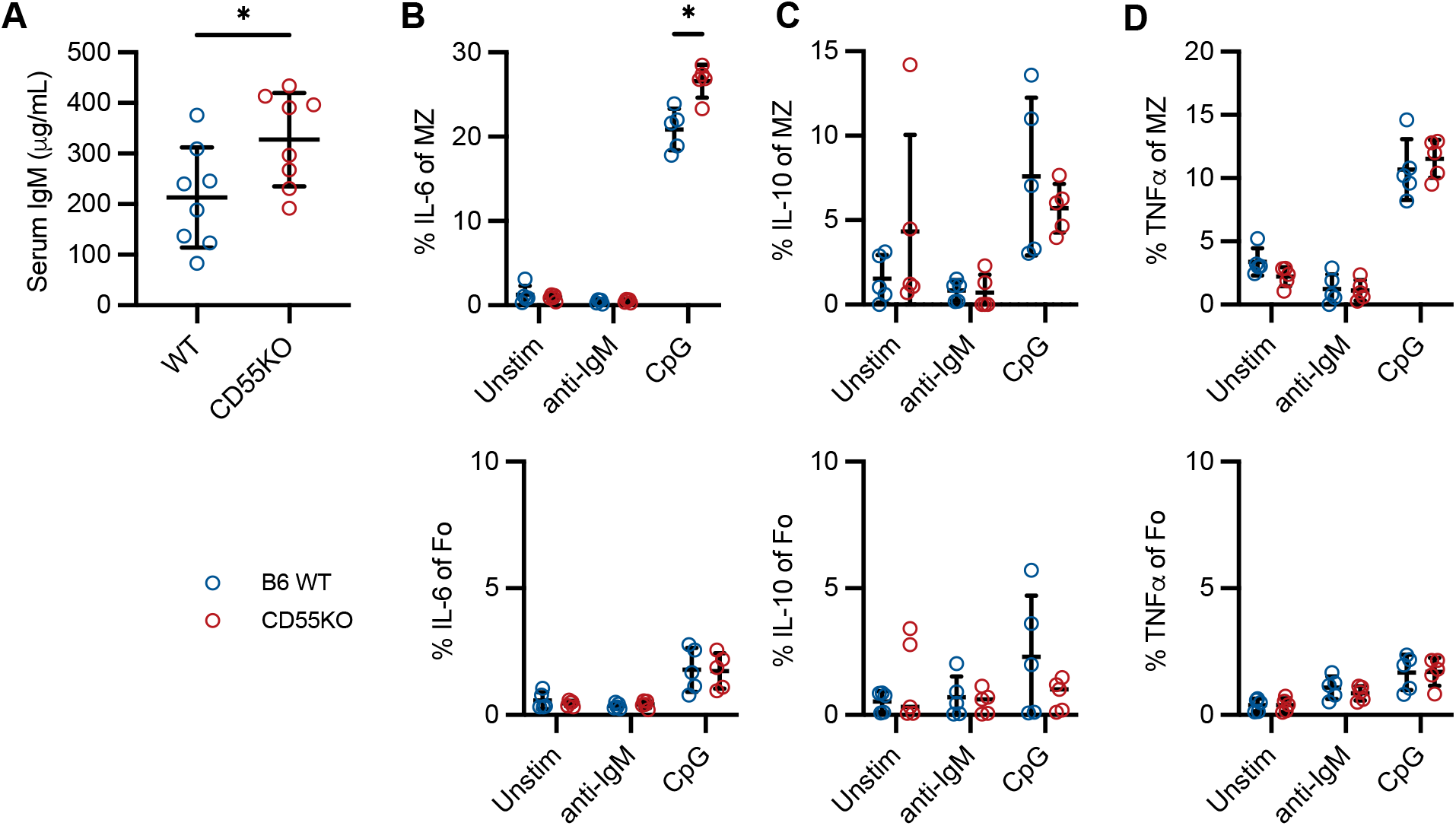
CD55 regulates baseline serum IgM level and the percentage of MZ B cells expressing IL-6. **A** Serum IgM levels in WT and CD55 KO mice. Percentage of MZ or follicular (Fo) B cells expressing **B** IL-6, **C** IL-10, or **D** TNFα from WT or CD55 KO mice after culturing in media alone or with anti-IgM or CpG. Statistical significance determined using t-test adjusted for multiplicity using Holm-Šídák’s correction. Error bars represent mean +/- standard deviation. All comparisons not significant except where marked. *p≤0.05.

In summary, we showed that CD55 regulates MZ B cell homeostasis and function. Loss of CD55 is not required for the MZ population to be established, but its loss prevents expansion of the MZ population through increased apoptosis. We also showed that CD55 regulates the expression of the cytokine IL-6 in MZ B cells. MZ B cells are strategically positioned in the marginal zone of the spleen where they are exposed directly to antigens in the blood and can trigger rapid immunoglobulin responses to antigens. MZ B cells also recognize self-antigen to aid in clearance of apoptotic cells, so their responses are necessarily highly regulated. Expansion of the MZ B cell population has been observed in multiple models of autoimmunity including transgenic strains producing B cells specific for autoantigens.^3, 4, 28, 29^ MZ B cells contribute to pathogenesis of disease in these models by secreting autoantibodies, producing cytokines, and presenting antigens to T cells.^4^

Our findings support a recent study that found that the interaction between CD55 on erythrocytes and CD97 on MZ B cells is required for retention of MZ B cells within the marginal zone of the spleen.^18^ The loss of the CD55-CD97 interaction seen in CD55 deficiency could be one of the factors contributing to CD55 KO MZ B cells undergoing apoptosis at homeostasis. We also note that CD55 is not required for the MZ B cell population to be established in the spleen, supporting previous studies that the MZ B cell population is heterogeneous.^30, 31^

While complement regulators are known to alter function within germinal center B cells, non-B cell antigen presenting cells, and T cells, this is the first report to our knowledge of a complement regulator altering MZ B cell function.^11, 15, 32-34^ Our findings augment those of a different complement regulator, complement factor H (CFH). CFH deficiency conferred expansion of the MZ population with increased IgM and IgG dsDNA titers.^35^ CFH is a fluid phase complement regulator that has decay acceleration properties like CD55, but also serves as a cofactor for the breakdown of the complement activation product C3b to iC3b and C3d, which are recognized by CD21. CFH-deficient MZ B cells increased CD21 expression whereas we observed decreased CD21/CD35 expression on CD55 KO MZ B cells. Loss of CD21 was previously shown to be associated with increased IL-6 and IL-10 production by MZ B cells as well as increased proliferation in both MZ and follicular B cells.^4^ We observed increased IL-6 but no change in IL-10 expression suggesting that decreased CD21 expression may mediate some of our findings. Thus, complement has important but complex roles in regulating MZ B cell homeostasis.

Located in the marginal zone of the spleen where arterial blood is filtered and scavenged for pathogens, MZ B cells are spatially and metabolically positioned to sense and mount rapid immunoglobulin responses against blood-borne antigens.^2, 36, 37^ Their responses are also highly regulated since MZ B cells harbor self-reactive B cell receptors.^36, 38^ In this study, we found that loss of CD55 was associated with decreased survival in MZ B cells at homeostasis. We further showed that loss of CD55 was associated with a proinflammatory phenotype with increased IL-6 expression in response to TLR9 stimulation. These findings suggest that in addition to homing, that CD55 regulates MZ B cell function in response to TLR9 stimulation. We are currently investigating this possibility further with cell-specific knockouts. These findings have implications for modulating the balance between protection against pathogens while maintaining tolerance in autoimmunity.

## Acknowledgements

We thank the Alvin J. Siteman Cancer Center at Washington University School of Medicine and Barnes-Jewish Hospital in St. Louis, MO. for the use of the Siteman Flow Cytometry Core Laboratory, which provided cell sorting service. We thank the Genome Technology Access Center at the McDonnell Genome Institute at Washington University School of Medicine for help with genomic analyses. These Centers are partially supported by NCI Cancer Center Support Grant #P30 CA91842 to the Siteman Cancer Center from the National Center for Research Resources (NCRR), a component of the National Institutes of Health (NIH), and NIH Roadmap for Medical Research. We lastly thank the UL1 TR002345 grant for training received by Dr. Iris Lee while performing these studies. This publication is solely the responsibility of the authors and does not necessarily represent the official view of NCRR or NIH.

## Funding

Iris Lee holds a Dean’s Scholars Award from the Washington University Division of Physician-Scientists, which is funded by a Burroughs Wellcome Fund Physician-Scientist Institutional Award. Peggy L. Kendall is supported by the Department of Medicine, VA Merit I01Bx002882, and R01DK084246.

## Notes

### Competing Interest Statement

The authors have declared no competing interest.

