## Supplemental Figures for "The complement regulator CD55 modulates TLR9 signaling and supports survival in marginal zone B cells"

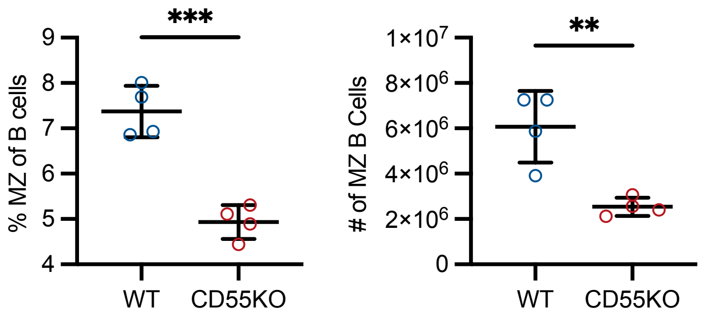


**Supplemental Figure 1. Differences in the MZ subset persist with age.** Percentage and number of MZ B cells in 17-week-old WT and CD55 KO mice. Statistical significance determined using Student's t-test. Error bars represent mean +/- standard deviation. **p≤0.01; ***p≤0.001

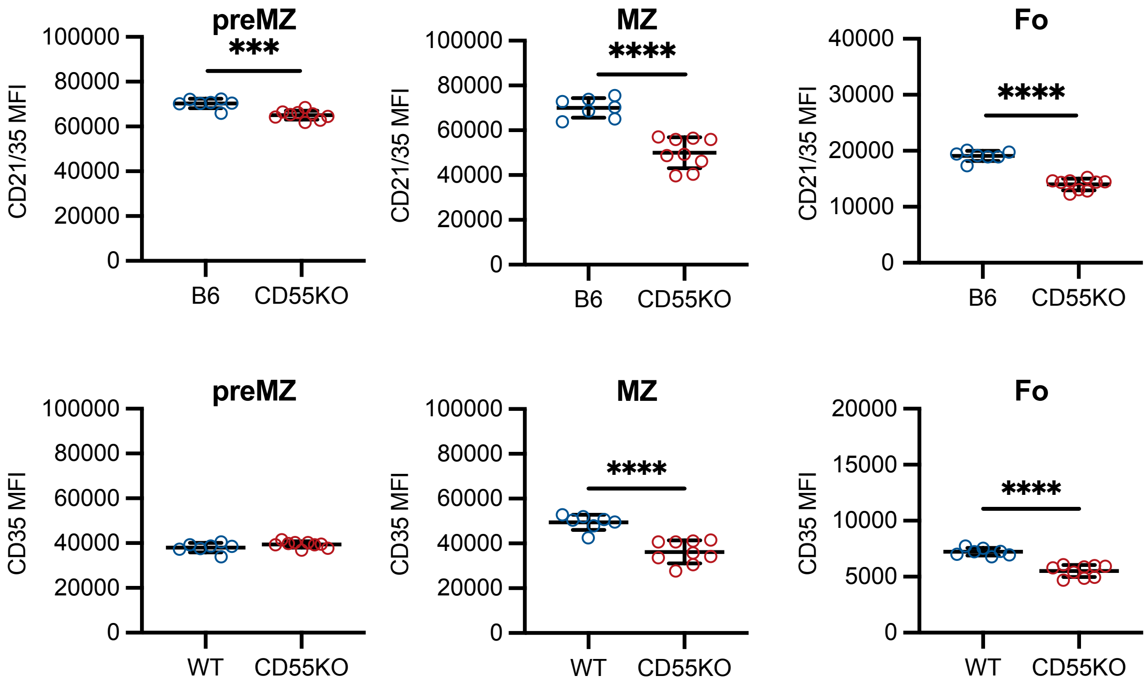


**Supplemental Figure 2. Loss of CD55 is associated with decreased CD21/CD35 in all B cell subsets.** CD21/35 and CD35 expression in preMZ, MZ, and follicular (Fo) B cells in 10- to

12-week-old mice. Statistical significance determined using Student's t-test. Error bars represent mean +/- standard deviation. ***$p \leq 0.001$; ****$p \leq 0.0001$.
